# A critical role of the T3SS effector EseJ in intracellular trafficking and replication of *Edwardsiella piscicida* in non-phagocytic cells

**DOI:** 10.1101/483032

**Authors:** Lingzhi Zhang, Jiatiao Jiang, Tianjian Hu, Jin Zhang, Xiaohong Liu, Dahai Yang, Yuanxing Zhang, Qin Liu

## Abstract

*Edwardsiella piscicida* (*E. piscicida*) is an intracellular pathogen within a broad spectrum of hosts. Essential to *E. piscicida* virulence is its ability to survive and replicate inside host cells, yet the underlying mechanisms and the nature of the replicative compartment remain unclear. Here, we characterized its intracellular lifestyle in non-phagocytic cells and showed that intracellular replication of *E. piscicida* in non-phagocytic cells is dependent on its type III secretion system. Following internalization, *E. piscicida* is contained in vacuoles that transiently mature into early endosomes, but subsequently bypasses the classical endosome pathway and fusion with lysosomes which depends on its T3SS. Following a rapid escape from the degradative pathway, *E. piscicida* was found to create a specialized replication-permissive niche characterized by endoplasmic reticulum (ER) markers. We also found that a T3SS effector EseJ is responsible for intracellular replication of *E. piscicida* by preventing endosome/lysosome fusion. Furthermore, *in vivo* experiments confirmed that EseJ is necessary for bacterial colonization of *E. piscicida* in both mice and zebrafish. Thus, this work elucidates the strategies used by *E. piscicida* to survive and proliferate within host non-phagocytic cells.

**Author summary:** *E. piscicida* is a facultative intracellular bacterium associated with septicemia and fatal infections in many animals, including fish and humans. However, little is known about its intracellular life, which is important for successful invasion of the host. The present study is the first comprehensive characterization of *E. piscicida*’s intracellular life-style in host cells. Upon internalization, *E. piscicida* is transiently contained in Rab5-positive vacuoles, but the pathogen prevents further endosome maturation and fusion with lysosomes by utilizing an T3SS effector EseJ. In addition, the bacterium creates an specialized replication niche for rapid growth via an interaction with the ER. Our study provides new insights into the strategies used by *E. piscicida* to successfully establishes an intracellular lifestyle that contributes to its survival and dissemination during infection.

## Introduction

Intracellular pathogens often invade host cells as a means of escaping extracellular immune defenses and creating a safe niche for replication. However, internalized pathogens are not entirely protected, as they are normally routed to lysosomes for degradation. Invasive pathogens must devise strategies to avoid this. Typically, intracellular pathogens either (i) reside within a customized, membrane-bound compartment, which limits trafficking along the endosomal pathway, as observed for *Legionella* [1], *Brucella subspp* [2] and *Salmonella* [3], or (ii) rupture and escape their vacuole to reside and replicate in the host cytosol, as in the case for *Shigella*, *Listeria*, and *Rickettsia subspp* [4].

Many pathogenic bacteria are found to proliferate in a membrane-bound compartment. These bacteria adopt different strategies to survive after phagocytosis. Some bacteria, such as *Salmonella* [5] and *Coxiella burnetiid* [6], survive and proliferate in an acidic compartment. Other pathogens avoid lysosomal fusion by blocking phagosome maturation, such as *Mycobacterium tuberculosis* [7], or by hijacking the eukaryotic secretory pathway, such as *Legionella pneumophila* [1]. *E. piscicida* was reported to reside within membrane-bound vacuoles (ECVs) after infection of both phagocytic and non-phagocytic cells [8,9]. However, the mechanism by which the bacterium evade lysosomal degradation remains unclear.

Host cell manipulation by pathogenic bacteria is largely mediated through the delivery of an arsenal of virulence proteins called effectors to the host cell cytosol [10,11]. *Legionella pneumophila* produce multiple effector proteins which specifically target host proteins such as Arf1, Rab1 and Sec22b to ultimately create a replicative organelle [12]. *Salmonella typhi* serovar Typhimurium is known to regulate *Salmonella*-containing vacuole (SCV) trafficking via the action of SPI-2 T3SS-delivered effectors [3]. For example, SifA targets the host GTPase Rab9 to inhibit the process of Rab9-dependent M6PR recycling [13] and SopD2 targets the host GTPase Rab7 to perturb endocytic trafficking [14]. Previous studies have shown that T3SS and T6SS mechanisms are essential for the virulence of *E. piscicida* [15]. An increasing number of T3SS and T6SS effectors have been identified, including EseG [16], EseJ [17], EseH [18], EseK [19] and EvpP [20]. EseG was reported to localize to the ECV membrane, but its function remains undefined [21]. EseJ was reported to be involved in the adhesion stage during infection [17]. However, the virulence factors involved in the regulation of replication of *E. piscicida* in host cells remain unknown.

Given previous findings supporting the ability of *E. piscicida* to invade, survive, and replicate within non-phagocytic cells [8], the goal of the present work was to uncover the strategies and molecular mechanisms used by this pathogen to circumvent lysosomal routing and establish a replicative niche within the host. We have identified an effector EseJ that is required for intracellular replication in a specialized vacuole that is important for *E. piscicida* replication inside host cells. We found that EseJ acts by inhibiting lysosome degradation of the pathogen which we find is important for systemic infection in vivo. Through these strategies, *E. piscicida* successfully establishes an intracellular lifestyle that could contribute to its survival and dissemination during infection.

## Results

### Intracellular replication of *E. piscicida* in non-phagocytic cells depends on its T3SS but not T6SS

*E. piscicida* prefers an intracellular lifestyle upon infection in either epithelial [8] or phagocytic cells [23]. However, the virulence factors involved in such an intracellular process remain undefined. Considering that T3SS and T6SS are the most important virulence factors for *E. piscicida*, we first monitored the survival and replication of both the wild-type EIB202 and the isogenic T3SS or T6SS mutant strains in three different non-phagocytic cells, HeLa, Caco-2 and ZF4. Both EIB202 and ΔT6SS infection of nonphagocytic cells, followed by gentamicin-induced death of extracellular bacteria, revealed a progressive increase in intracellular bacterial numbers over time (Fig 1A). In contrast, no replication was observed in the ΔT3SS mutant, indicating that a functional T3SS, but not a T6SS, is required for intracellular survival of *E. piscicida* in non-phagocytic cells. In order to visualize bacterial invasion and intracellular replication, HeLa cells were infected with green fluorescent protein (GFP)-tagged *E. piscicida* strains, confocal microscopy images were acquired and the number of intracellular bacteria in EIB202-infected cells were scored over time. About 15% of HeLa cells containing hyper-replicating bacteria were observed after EIB202 and ΔT6SS infection for 8 h, but not with the ΔT3SS mutant strain (Fig 1B and 1C). Collectively, these results suggest that *E. piscicida* can survive and replicate in non-phagocytic cells via a mechanism mediated by the T3SS, but not T6SS.

**Fig 1.**
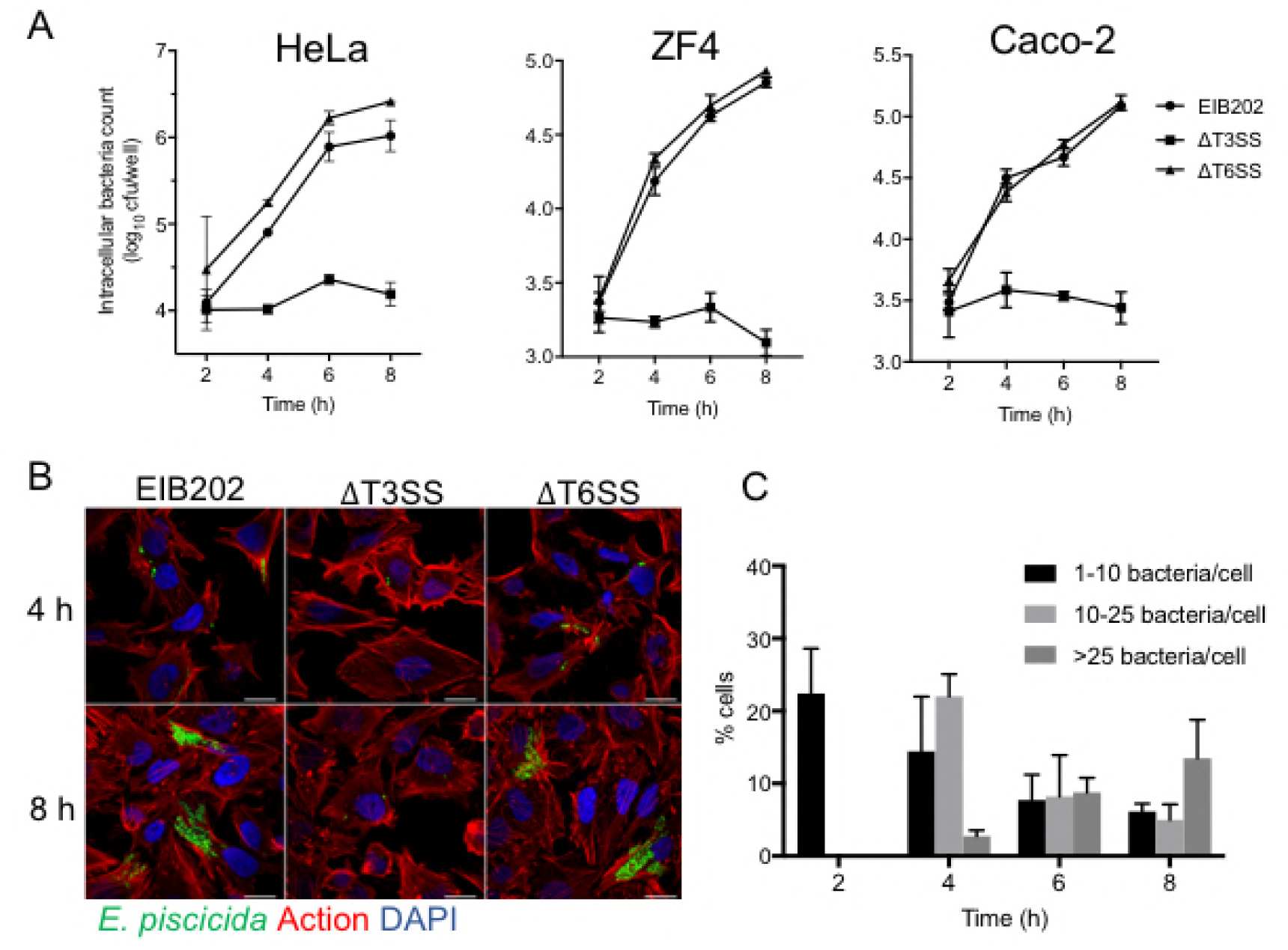
*E. piscicida* replicates in non-phagocytotic cells dependent on T3SS but not T6SS. (A) HeLa cells, ZF4 and Caco-2 cells were infected with *E. piscicida* EIB202, ΔT3SS or ΔT6SS at an MOI of 100 for 1 h, followed by treatment with 100 μg/ml gentamicin for 1 h to kill extracellular bacteria. Intracellular bacteria at different time point were quantified by lysis, serial dilution and viable counting on TSB agar plates. (B) Confocal microscopy of HeLa cells infected with GFP-labeled *E. piscicida* EIB202, ΔT3SS or ΔT6SS at 4 and 8 h. Data are representative of at least three experiments, and representative microscopic images are shown. Filamentous actin was stained by rhodamine-phalloidin (red), and DNA was stained by DAPI (blue). Scale bars, 20 μm.(C) Percentage of infected cells containing one to ten, ten to twenty-five or more than twenty-five intracellular wild-type *E. piscicida* over time.

### *E. piscicida* prevents endosome maturation and lysosome fusion

Once inside host cells, invasive bacteria either replicate within the endosome or escape the vacuole and replicate in the cytoplasm. Consistent with our previous study [21], *E. piscicida* located within vacuolar compartment (s) after infection which we named the *E. piscicida*-contained vacuoles (ECVs) (S1A Fig). Galectin-3 is a β-galactoside binding protein that is specifically recruited to disrupted pathogen-containing vacuoles [24]. To further investigate if *E. piscicida* remains inside the vacuoles during its whole intracellular lifecycle, the presence of galectin-3 around the ECV was assessed using fluorescence microscopy. Less than 10% of vacuoles harboring WT *E. piscicida* co-localized with galectin-3 in HeLa cells over time (S1B Fig). These data suggest that *E. piscicida* EIB202 resides and replicates inside pathogen-containing vacuoles throughout the course of infection.

Next, we wished to understand the strategies used by this pathogen to establish and maintain its intracellular life cycle. Following internalization, foreign particles and many bacteria are usually found within membrane-bound compartments that sequentially develop into early and late endosomes for ultimate fusion with lysosomes, where the particles are degraded. We monitored the acquisition of endosomal markers and lysosomal fusion in *E. piscicida*-containing vesicles over time using confocal microscopy. Early after invasion (1 h after infection), over 70% of both intracellular wild-type and ΔT3SS mutant were found enclosed within vacuoles that co-localized with the early endosomal protein Rab5 (Fig 2A and 2B), indicating interactions with early endosomes. These interactions were transient, as Rab5 colocalization rapidly decreased to 5% and 3% by 4 h post-gentamicin incubation after infection with both wild-type EIB202 and the ΔT3SS mutant strains (Fig 2A and B). As Rab5 colocalization was progressively lost, an increasing number of ΔT3SS mutant bacteria colocalized with the late endosomal markers Rab7 and lysosome-associated membrane protein 1 (Lamp-1) over time, which is consistent with vacuolar maturation (Fig 2B-E). In contrast, the majority of the vacuoles containing wild-type EIB202 were negative for both Rab7 and Lamp1 (Fig 2B-E). These results suggest wild-type *E. piscicida* transiently interacts with early endosomes, but avoids endosome maturation during infection.

**Fig 2.**
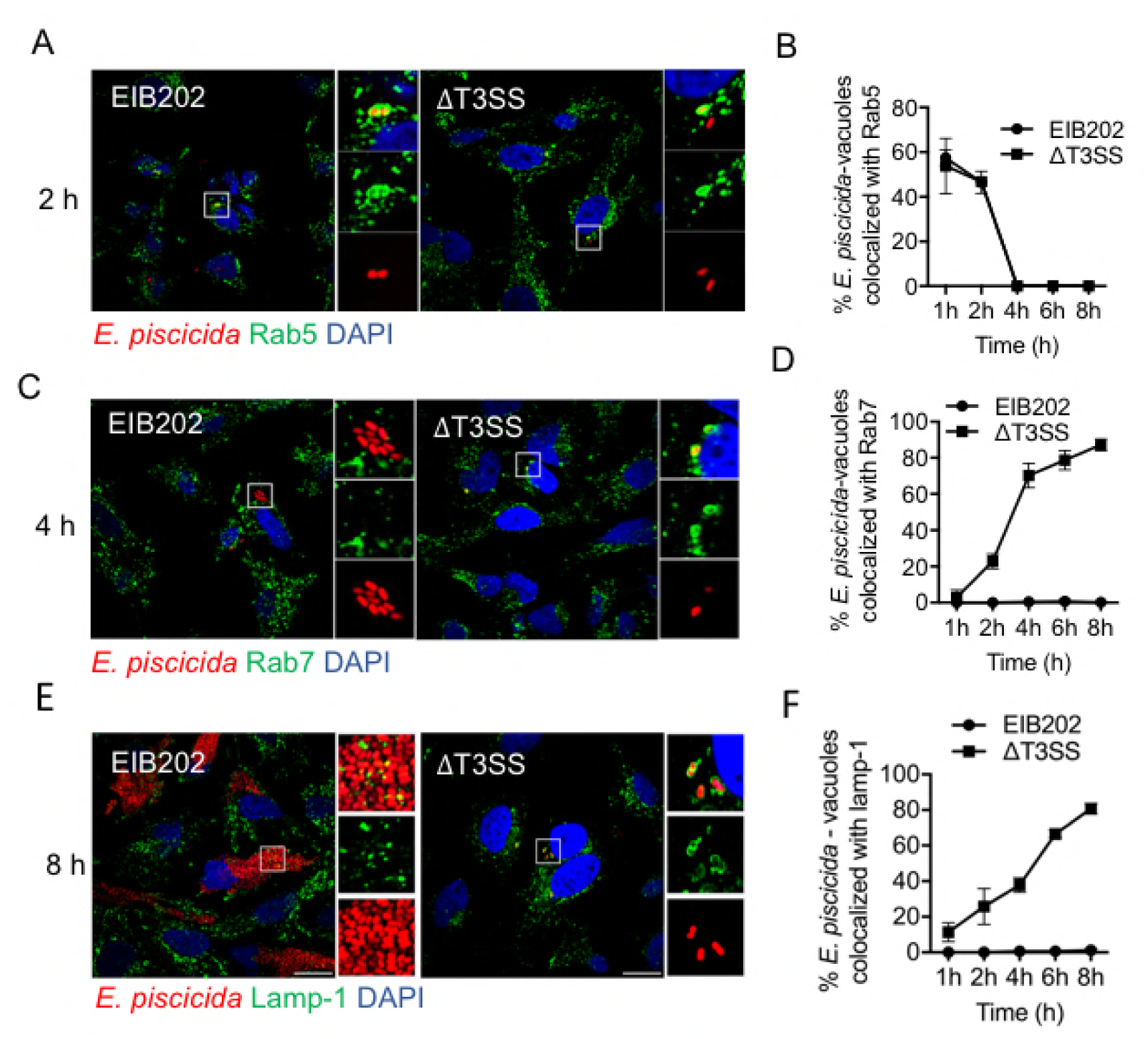
*E. piscicida*-contained vacuoles evades maturation into late endosomes. (A, C and E) Representative confocal micrographs of HeLa cells infected with RFP-expressing *E. piscicida* wild-type EIB202 or ΔT3SS and incubated with gentamicin for the times indicated on the sides of the panels. Cells were pre-transfected with GFP-Rab5(A [Green]), GFP-Rab7 (C [Green]) or immunostained for lamp-1 (E [Green]). DNA was stained using DAPI (blue). White boxes indicate the magnified area to the right of each panel. Scale bars, 20 μm. (B, D and F) Quantifications of ECVs colocalization with Rab5 (B), Rab7 (D) and lamp-1 (F) for the indicated times. Over 30 cells were analyzed for each condition. Values are means ± SD (n= 3).

Considering that luminal acidification is another critical characteristic of endosome maturation, we used the fixable acid tropic probe LysoTracker to monitor acidic organelles in infected cells. A major overlap was found between the dye and ΔT3SS mutant strain, but not wild-type EIB202 as early as 2 h (Fig 3A and B). Thus, these results suggest the ECVs formed by the wild-type bacterium avoid vacuolar acidification and maturation by perturbing the fusion with lysosomes. To further understand this process, we assessed ECV co-localization with TR-dextran. Prior to bacterial infection, cells were pulsed with TR-dextran for 6 h followed by overnight chase in dye-free medium to ensure that the probe is delivered from early and recycling endosomes to lysosomes [25]. Confocal immunofluorescence and quantification data showed that the majority of wild-type EIB202-containing vacuoles did not co-localize with TR-dextran (Fig 3C and D). In contrast, when cells were infected with the ΔT3SS strain, more than 70% of the ECVs did co-localize with TR-dextran at 8 h post-infection (Fig 3C and D). Taken together, these results indicate that *E. piscicida* utilize its T3SS to successfully evade lysosomal fusion and ultimately replicate in nonacidic compartments lacking lysosomal or late endosomal characteristics.

**Fig 3.**
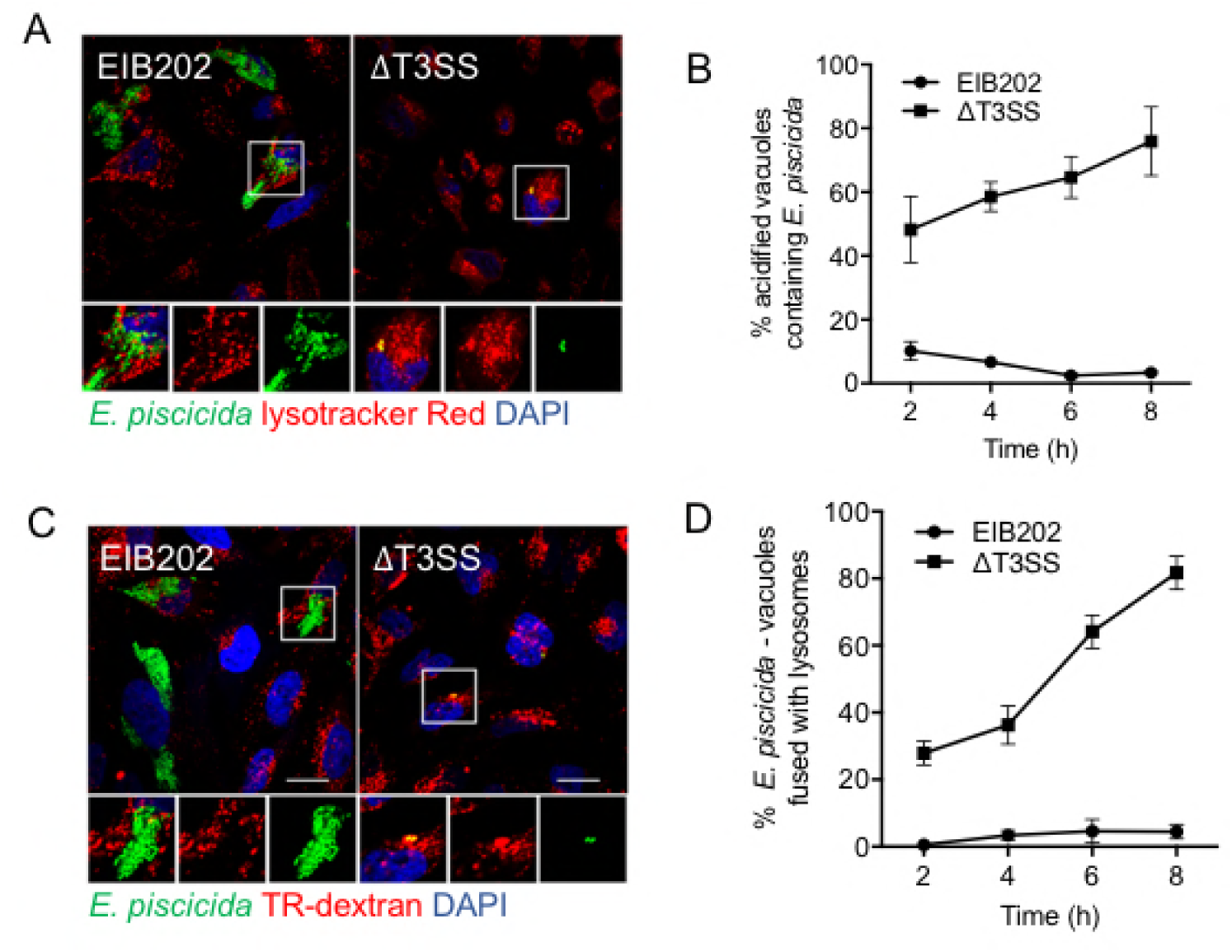
*E. piscicida*-contained vacuoles evades fusion with lysosomes by T3SS. (A) Representative confocal micrograph of HeLa cells infected with GFP-labeled *E. piscicida* EIB202 or ΔT3SS for 1 h and incubated with 100 μg/ml gentamicin for 8 h. During the last 30 min of antibiotic treatment, samples were added with 75 nM Lysotracker Red DND-99 (red). DNA was stained using DAPI (blue). White boxes indicate the magnified area to the below of each panel. Scale bars, 20 μm. (B) Quantification of acidified vacuoles containing *E. piscicida* EIB202 or ΔT3SS at the indicated gentamicin incubation times. Values are means±SD from over 30 cells (n= 3). (C) Representative confocal micrograph of HeLa cells preloaded with 1 mg/ml Texas Red dextran (red) for 6 h and chased overnight, after which cells were infected with GFP-labeled *E. piscicida* EIB202 or ΔT3SS for 1 h and incubated with gentamicin for 8 h. Scale bars, 20 μm. (D) Quantification of *E. piscicida*-contained vacuoles colocalizied with Dextran at the indicated gentamicin incubation times. Values are means±SD from over 30 cells (n=3).

### ECVs acquire ER markers during maturation into a replicative organelle

Our results iundicate that wild-type *E. piscicida* circumvented the classical endocytic pathway to establish a specialized replication-permissive niche. This raises the possibility that the bacterium may interact with other intracellular compartments. The endoplasmic reticulum (ER) was reported to be recruited and hijacked by many intracellular pathogens to create their replication niche [1,26]. We first used a specific ER-tracker dye to assess the intracellular ER structures and analyzed the presence of ER marker in bacterium-containing vacuoles. Surprisingly, ER-tracker labeling was present in the EIB202 replication compartment located at the perinuclear regions of HeLa cells, but this was excluded in ΔT3SS-containing vacuoles (Fig 4A and B). The same result was observed when we studied the distribution of the ER membrane-bound lectin calnexin in EIB202 or ΔT3SS-infected cells (Fig 4C and D). Together, these data suggest that *E. piscicida* replicates inside ER-enriched vacuoles.

**Fig 4.**
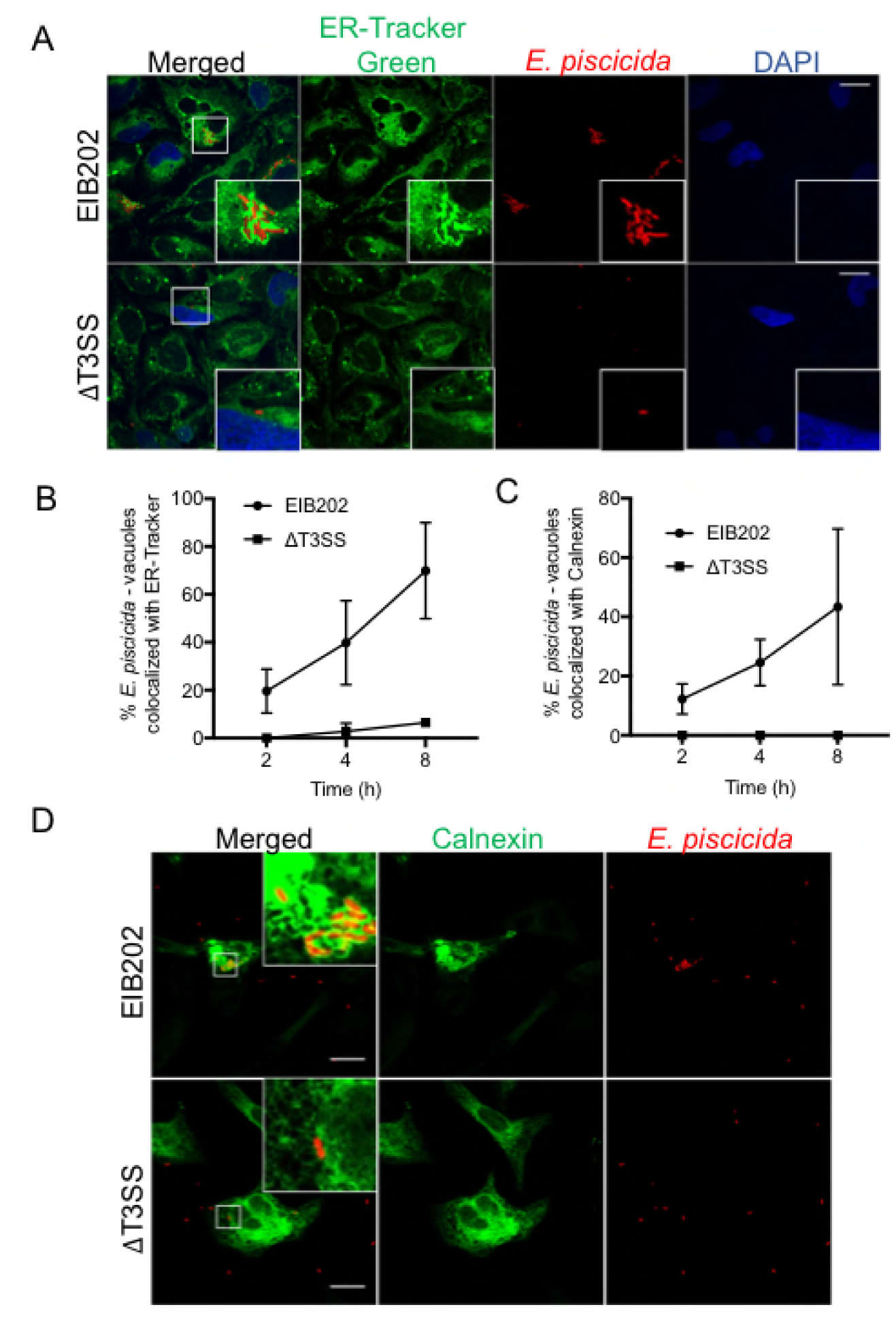
*E. piscicida* resides and replicates in ER-characterized vacuoles. (A) Representative confocal micrograph of HeLa cells infected with RFP-labeled *E. piscicida* EIB202 or ΔT3SS for 1 h and incubated with 100 μg/ml gentamicin for 4 h. During the last 30 min of antibiotic treatment, cells were washed with HBSS and stained with 100 nM ER-tracker (green). DNA was stained using DAPI (blue). Insets are enlarged from the indicated area. Scale bars, 20 μm. (B) and (C) Quantifications of *E. piscicida*-contained vacuoles colocalization with ER-tracker (B) or Calnexin (C) at the indicated gentamicin incubation times. Values are means±SD from over 30 cells (*n*=3).(D) Representative confocal micrograph of HeLa cells infected with RFP-labeled *E. piscicida* EIB202 or ΔT3SS for 1 h and incubated with 100 μg/ml gentamicin for 4 h. Cells were immunostained with Calnexin (GFP). DNA was stained using DAPI (blue). Insets are enlarged from the indicated area. Scale bars, 20 μm.

### A T3SS effector EseJ is responsible for *E. piscicida*’s intracellular replication

The finding that the T3SS plays a critical role in the inhibition of endosome maturation and *E. piscicida* lysosome degradation raised the question of what T3SS effectors are involved in this process. To date, several *E. piscicida* T3SS effectors including EseG [16], EseJ [17], EseH [18] and EseK [19] have been identified. To assess the role of individual effectors, we tested the ability of WT and isogenic *E. piscicida* effector mutants to replicate inside non-phagocytic host cells. We found that only the *eseJ* mutant showed a marked deficiency in intracellular replication compared to wild-type bacteria as assessed by CFU intracellular counts (Fig. 5A). To determine whether the impaired ability of the *eseJ* mutant to replicate intracellularly was attributed to effects on lysosome fusion and degradation, we investigated the characteristics of the Δ*eseJ*-containing compartments over time. Coincident with the intracellular fate of ΔT3SS, Δ*eseJ*-containing vacuoles progressively co-localized with Rab7 (S2A Fig) and Lamp-1(Fig 5B and S2B Fig). Moreover, the mature Lamp-1-positive Δ*eseJ*-containing vacuoles were found to be fused with lysosomes as assessed using the acidification probe LysoTracker Red DND-99 and pre-loaded Dextran (Fig 5C and 5D). These findings suggest that the effector EseJ is critical for *E. piscicida’*s replication inside cells by disrupting vacuolar trafficking to the lysosome during infection.

**Fig 5.**
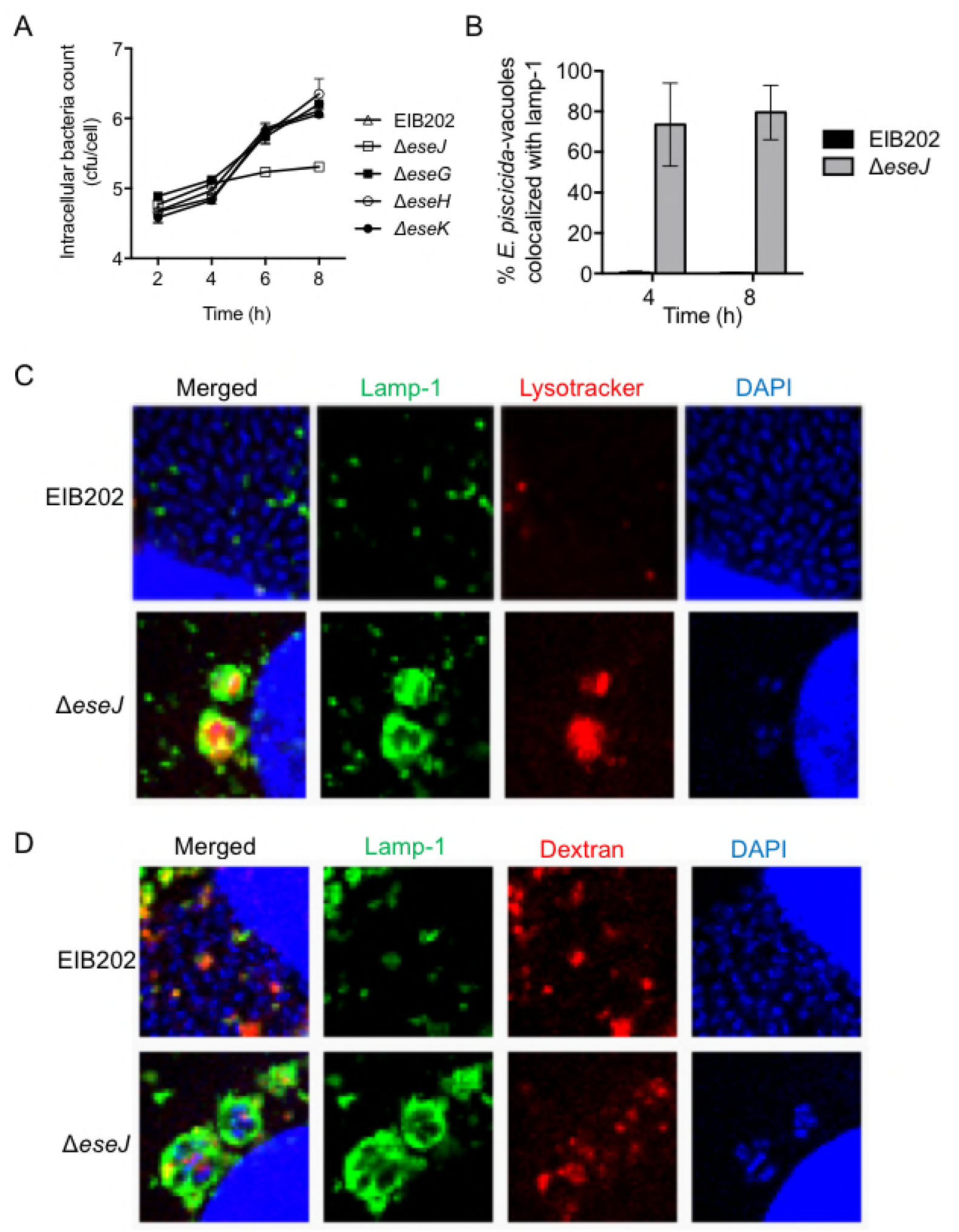
The T3SS effector EseJ is critical for intracellular replication of *E. piscicida* in HeLa cells. (A) HeLa cells were infected with *E. piscicida* EIB202, Δ*eseJ*, Δ*essG*, Δ*essH* or Δ*eseK* at an MOI of 100 for 1 h, followed by treatment with 100 μg/ml gentamicin for 1 h to kill extracellular bacteria. Intracellular bacteria at different time point were quantified by lysis, serial dilution and viable counting on TSB agar plates.(B) Quantifications of ECVs colocalization with lamp-1 at the indicated gentamicin incubation times. Values are means±SD from over 30 cells (n=3). (C-D) Representative confocal micrograph of HeLa cells infected with GFP-labeled *E. piscicida* EIB202 or Δ*eseJ* for 1 h and incubated with 100 μg/ml gentamicin for 8 h. Cells were stained with 75 nM Lysotracker Red DND-99 (C, red) or preloaded with1 mg/ml Texas Red dextran (red) for 6 h and chased overnight. DNA was stained using DAPI (blue) and late endosomes/lysosome were immunostained with lamp-1. Scale bars, 20 μm.

To further assess the function of EseJ, we analyzed the effect of EseJ expression on the transport and the degradation of exogenously added DQ-Red bovine serum albumin (BSA), which emits red fluorescence upon proteolytic degradation and is used as a sensitive indicator of lysosomal activity [14]. Bright punctate signal intensity of DQ-Red BSA were significantly attenuated in cells stably expressing EseJ compared with that observed in cells expressing vector alone (Fig 6A and 6B), indicating that EseJ expression suppresses lysosome function. Consistently, cells infected with wild-type, but not ΔT3SS or Δ*eseJ E. piscicida* mutants displayed a remarkable decrease in DQ-Red BSA fluorescence intensity (S2C Fig). Next, we investigated the delivery of endosomal cargo to lysosomes by pre-loading cells with dextran 488 prior to transfection and then treated the cells with rhodamine dextran. In line with the results shown above, the dextran derivatives co-localized with a Mander’s coefficient of more than 0.5 in control cells, suggesting significant endosome-lysosome fusion, whereas EseJ-HA expression resulted in significantly less co-localization (Fig 6 C and 6D). Collectively, these results demonstrate that T3SS effector EseJ is both necessary and sufficient to block endocytic trafficking to lysosomes and consequently critical for *E. piscicida*’s intracellular replication.

**Fig 6.**
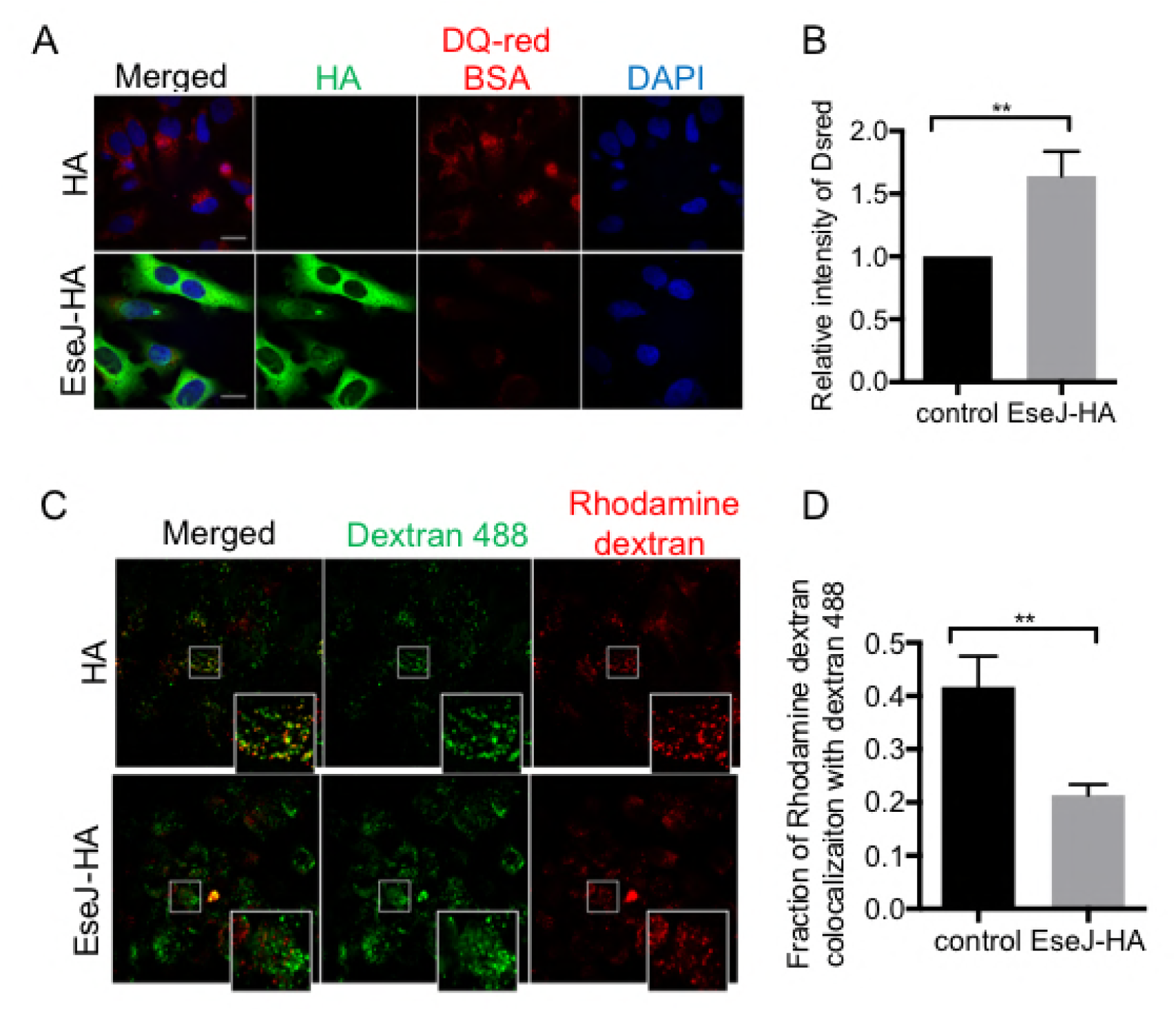
The T3SS effector EseJ is necessary and sufficient to block endocytic trafficking to lysosomes. (A) HeLa cells which stably expressed HA tagged EseJ or HA tag only were incubated with DQ-Red BSA (red) for 1 hr. Images were acquired after an additional 4 hr chase. (B) Quantification of DQ-Red BSA signal for (A), ** *p*<0.01. (C) HeLa cells which preloaded with dextran 488 for 8 hr, transfected with HA or EseJ-HA (green), and then pulsed with rhodamine dextran for 30 min. Imaging was performed after 2 hr of chase. (D) Quantification of colocalization signal between two dextran derivatives for (C), ** *p*<0.01.

### Role of EseJ in *E. piscicida*’s infection in vivo

*E. piscicida* T3SS was reported to act as a critical virulence factor in disease pathogenesis in both mouse and fish infection models [23, 27]. To assess the role of the T3SS effector EseJ in animal infection, C57BL/6 wild-type mice were orally infected with wild-type and isogenic Δ*eseJ E. piscicida* strains. Compared to wild-type EIB202, the Δ*eseJ* strain showed reduced bacterial burdens in the cecum and intestinal lumen as well as systemic sites including the liver, spleen and kidneys (Fig 7A). Likewise, reduced colonization of the Δ*eseJ* mutant was observed in zebrafish larvae after infection when compared to the wild-type bacterium (Fig 7B). Notably, zebrafish infected with wild-type *E. piscicida* showed marked mortality, with ~75% of the animals succumbing by day 3–4 post-infection whereas only ~50% of fish succumbed to infection with the *eseJ* mutant (Fig. 7C). Collectively, these data suggest that the effector EseJ contributes to *E. piscicida* colonization and virulence in vivo.

**Fig 7.**
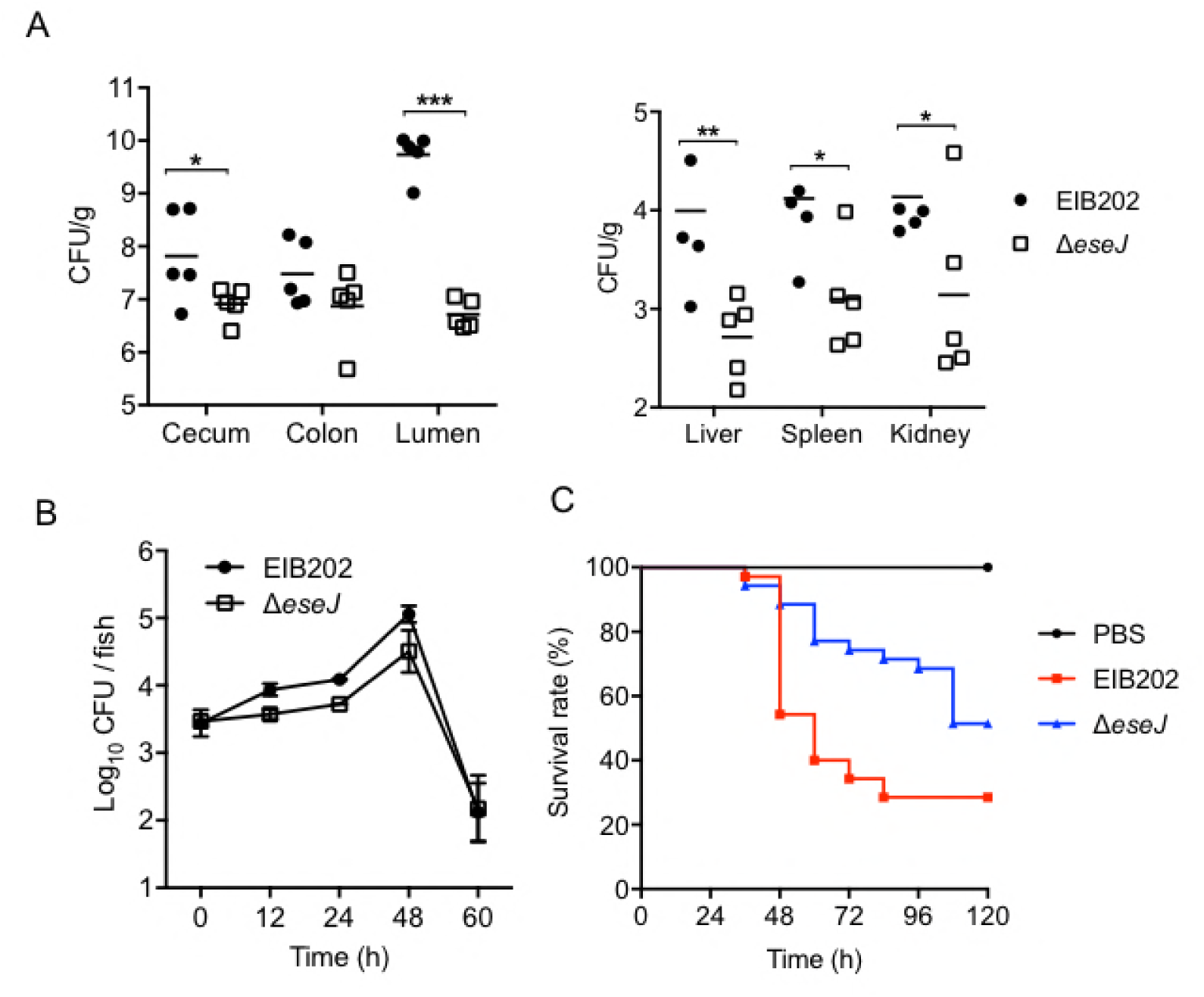
Critical role of EseJ in promoting virulence and colonization of *E. piscicida in vivo*. (A) Bacteria burden in the colon, cecum, lumen, liver, spleen, and kidney of mice was measured after orally-infection with EIB202 or Δ*eseJ* (2.5 × 10^7^ cfu/g) at 48 hpi. * p < 0.05, ** *p* < 0.01, *** *p* < 0.001. (B) Bacteria burden in zebrafish larvae was measured at indicated time points after infection by immersion with 1 × 10^5^ cfu/ml EIB202 or Δ*eseJ*. N = 5 fish per group per time point. Data are representative of at least three experiments. (C) Survival of zebrafish infected with EIB202 or Δ*eseJ* (50 cfu/fish). N = 35 fish per group. Data shown are from at least three representative experiments.

## Discussion

*E. piscicida* was recently shown to invade and proliferate within many non-phagocytic cells [8], but the mechanism by which the bacterium its own survival inside host cells remained unclear. Here we report a comprehensive description of how *E. piscicida* turns the intracellular environment into a hospitable niche that allows for efficient bacterial replication. Subversion of the phagocytic pathway by intracellular bacteria is a general mechanism to establish an appropriate replication niche. Pathogens are known to adopt diverse strategies to disrupt the maturation process at different stages and to prevent its delivery into a phagolysosome. For example, *Mycobacterium* remains within an early endosomal compartment [7] that excludes the vacuolar ATPase, thus inhibiting the acidification of the bacterial phagosome. The maturation of the SCV is arrested at a late endosome-like stage, selectively excluding proteins such as mannose 6-phosphate receptors (MPR) and lysosomal cathepsin proteins [13]. In the present study, we tracked the acquisition of endosomal markers and lysosomal fusion in *E. piscicida*-containing vesicles over time using confocal microscopy and demonstrated that *E. piscicida* bypassed the classical endosome pathway after transiently interactions with early endosomes. Our studies indicate that using this strategy, *E. piscicida* disrupts endosomal maturation and evades lysosome degradation.

Our study characterized an important contribution of the T3SS effector EseJ in regulating endocytic trafficking of *E. piscicida* within host cells. Intracellular expression of EseJ was found necessary and sufficient to block endocytic progression to lysosomes (Fig 5 and Fig 6). Notably, the *eseJ* mutant was greatly impaired in intracellular replication when compared to the wild-type bacterium, indicating that EseJ is an important factor for intracellular survival and replication of *E. piscicida*. However, it remais unclear how EseJ evades fusion with lysosomes to evade degradation. One possibility is that EseJ interacts with host small guanine nucleotide binding proteins (GTPases), phospholipids or other host proteins that are enriched and central for endocytic trafficking. The strategy of interacting with endosome-bounded proteins is an efficient tactic used by other pathogens to combat the host’s bactericidal defenses. For example, Mycobacterium tuberculosis (Mtb) secretes SapM, a phosphatase that removes PI(3)P from Mtb-containing vacuoles by converting it to PI, thereby arresting endosomal maturation [28]. *Legionella pneumophila* secreted VipD to interact with early endosomal protein Rab5 to protect from endosomal fusion [29]. Another question is whether EseJ act in concert with other virulence factors involved in the regulation of ECV trafficking which needs to be investigated in future studies.

Orchestration with other intracellular compartments and routing into a specialized compartment favorable for replication is another important mechanism for the survival of bacterial pathogens inside host cells. For example, biogenesis of *Legionella*-replicative compartments depends upon a rapid interception of COPI-dependent vesicular trafficking from endoplasmic ER exit sites [12]. Formation of SCVs is associated with the Golgi apparatus and induces endosomal tubulations that extend towards the cell periphery [30]. Interestingly, we observed obvious ER characteristic associated with ECVs. However, how *E. piscicida* recruits and interacts with ER remains to be elucidated.

Overall, our studies demonstrate a complex and deliberate intracellular life cycle of *E. piscicida* in non-phagocytic cells (see model in Fig. 8). The bacterium not not only invades the host cells, but also subverts trafficking of bacterium-containing vacuoles through the endosomal pathway and translocation to an specialized aggressively replication niche. Moreover, we showed that a T3SS effector EseJ is essential for the intracellular replication by disrupting endosomal maturation and lysosome fusion, which is critical for virulence of *E. piscicida in vivo*.

**Fig 8.**
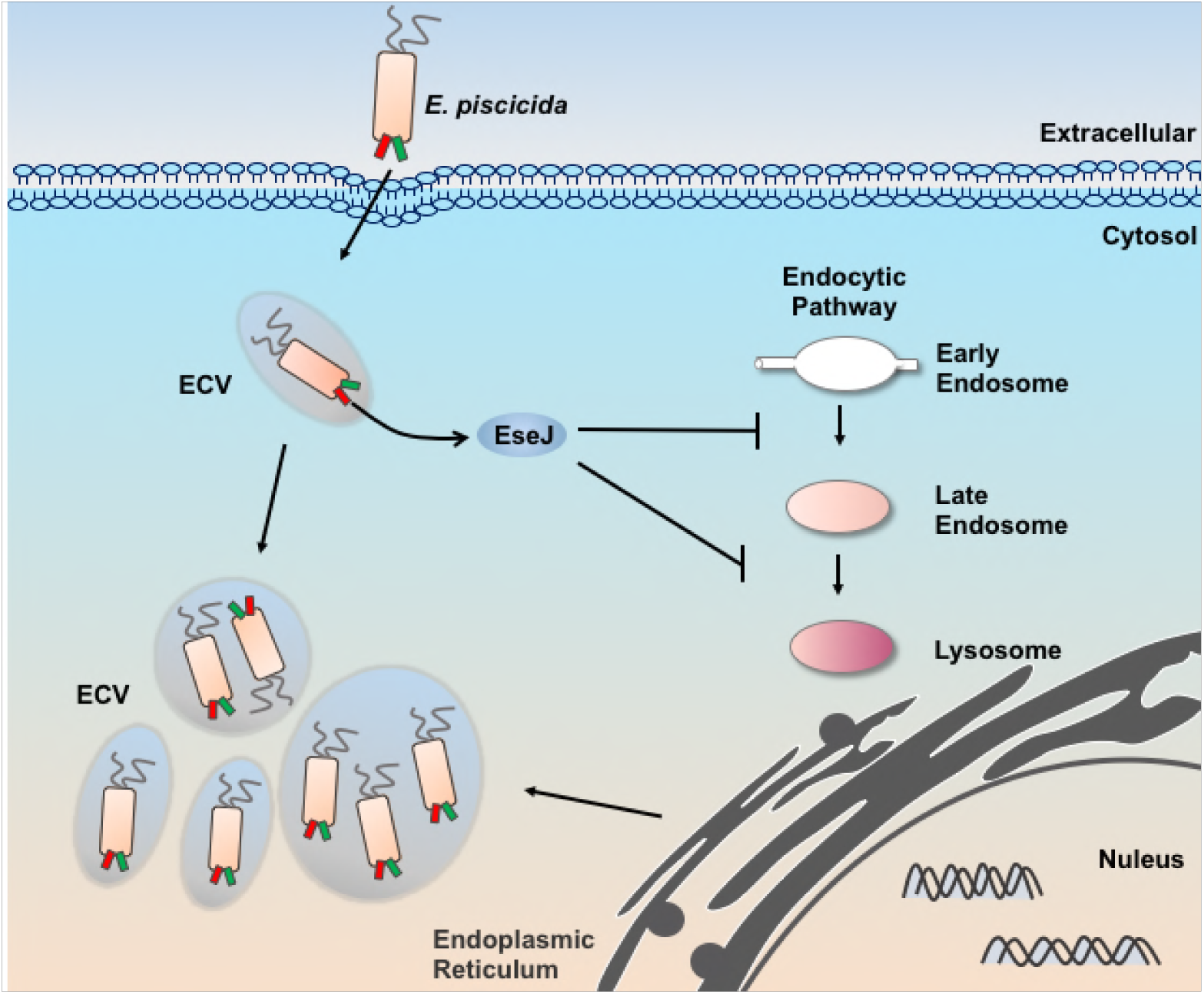
Proposed model for *E. piscicida* intracellular life cycle in non-phagocytic cells. After entry, intracellular wild type *E. piscicida* resides within vacuoles (ECVs) that interact with early endosomes. At intermediate stages of infection, an T3SS effector EseJ was secreted into cell cytosol to globally disrupt endocytic trafficking to lysosomes, which helps to protect ECVs from lysosome fusion. Therefore, these early ECVs bypass the regular lysosomal routing but contact with the ER to facilitate its later robust replication.

## Methods

### Ethics Statemen

The animal trials in this study were performed according to the Chinese Regulations of Laboratory Animals—The Guidelines for the Care of Laboratory Animals (Ministry of Science and Technology of People’s Republic of China) and Laboratory Animal-Requirements of Environment and Housing Facilities (GB 14925-2010, National Laboratory Animal Standardization Technical Committee). The license number associated with their research protocol was 20170912-08, which was approved by The Laboratory Animal Ethical Committee of East China University of Science and Technology. All surgery was performed under carbon dioxide anesthesia, and all efforts were made to minimize suffering.

### Bacterial strains and cell culture

Wild type *Edwardsiella piscicida* EIB202, the T3SS mutant and the T6SS mutant were constructed and grown as described previously [22]. For constitutive expression of GFP or mCherry, *E. piscicida* strains were electroporated with pUTt0456GFP or pUTt0456mCherry, respectively. HeLa cells (ATCC number CCL-2), Caco-2 cells (ATCC number HTB-37) and ZF4 cells (ATCC number CRL-2050) were all from China Center for Type Culture Collection. HeLa cells and Caco-2 cells were cultured at 37°C under 5% CO_2_ atmosphere in Dulbecco’s minimal Eagle’s medium (DMEM) supplemented with 10% fetal bovine serum (FBS), called growth medium (GM). ZF4 cells were cultured at 30°C under 5% CO_2_ atmosphere in GM.

### Construction of mutant strains

In-frame deletion mutants of the effector genes including *eseG*, *eseJ*, *eseH* and *eseK* were generated by the sacB-based allelic exchange as previously described. The fragments upstream and downstream of each effector gene were fused by overlap PCR. These fragments were then cloned into the sacB suicide vector pDMK and linearized with BglII and SphI, and the resulting plasmids were transformed into *Escherichia coli* (*E. coli*) CC118 λpir. The correct plasmids were then transformed into *E. coli* SM10 λpir and then conjugated into EIB202. The trans-conjugants with the plasmids integrated into the chromosome by homologous recombination were selected on tryptic soy agar (TSA) medium containing kanamycin (Km, 50 mg/ml) or colistin (Col, 12.5 mg/ml). To complete the allelic exchange for in-frame deletions, double-crossover events were counter-selected on TSA plates containing 10% sucrose. All of the mutants were confirmed by PCR amplification of the respective DNA loci, and subsequent DNA sequencing of each PCR product.

### Infection protocol

HeLa cells, Caco-2 cells or ZF4 cells were infected with *E. piscicida* strains at a multiplicity of infection (MOI) of 100. *E. piscicida* was grown overnight in tryptic soy broth (TSB) at 30°C with shaking, then diluted into fresh DMEM with standing at 30°C until OD_600_ reached 0.8. Harvested bacteria in phosphate-buffer saline (PBS) suspensions were added to cells according to MOI. To synchronize infection, plates were then centrifuged at 600 *g* for 10 min. At 1 h after incubation, cells were washed three times with PBS and then incubated with growth medium containing 100 μg/ml gentamicin for 1h to kill the extracellular bacteria, after which the gentamicin concentration was decreased to 10 μg/ml for the remainder of the experiment.

### Gentamicin protection assay

For enumeration of viable intracellular bacteria, bacteria were added to triplicate wells of HeLa cell monolayers for infection as described above. At each indicated time point, extracellular bacteria were killed with gentamicin. Monolayers were washed with PBS, and cells were lysed by incubation with PBS containing 1% Triton X-100 for 30 min at room temperature. The lysate was serially diluted in PBS and plated onto TSB agar plates. Plates were incubated at 30°C overnight for subsequent CFU enumeration.

### Labeling of subcellular compartments with dyes

For the acidification studies, 75 nM LysoTracker Red DND-99 (Invitrogen) was added to the samples 30 min prior to cell fixation. For labeling of lysosomes with Texas Red dextran, HeLa cells were treated with 100 μg/ml of Texas Red dextran (Invitrogen) for 12 h and chased overnight. For ER staining, cells were washed with HBSS and stained with 100 nM ER-tracker (green) for 30 min at indicated time point. For DQ Red BSA assay, after 8 h of infection or cells transfection, cells were incubated for 1 h in growth medium containing DQ Red BSA (0.25 mg/ml), washed with PBS, and incubated in growth medium for 4 h.

### Dextran 488 Loading and Rhodamine Dextran Pulse-chase

HeLa cells were seeded on coverslips in 24-well tissue culture plates at 2×10^5^ cells/well then incubated in presence of dextran Alexa Fluor® 488 (0.1 mg/ml) for 8 h. Cells were then washed twice with PBS, incubated with growth medium and transfected with vector or EseJ-HA overnight. The following day cells were incubated for 30 min in the presence of tetramethylrhodamine dextran (0.2 mg/ml), then washed twice with PBS, and the dye was chased for 2 h in regular growth medium.

### Immunofiuorescence and confocal microscopy

HeLa cells were seeded onto 24-well plates containing sterile coverslips at a density of 2×10^5^ cells/ml. Following infection with *E. piscicida* strains and gentamicin incubation for the indicated time, cells were washed with phosphate-buffered saline (PBS) and then fixed in 4% (v/v) paraformaldehyde for 10 min at room temperature. After washing with PBS, cells were blocked and permeabilized in PBS containing 10% (v/v) normal goat serum (NGS) and 1% (v/v) bovine serum albumin (BSA) and 0.1% (w/v) saponin (SS-PBS) for 10 min at room temperature. Primary antibody of LAMP-1(clone H4A3) and secondary antibodies were diluted in SS-PBS at appropriate dilutions and incubated serially for 1 h at room temperature. Between antibody incubations, coverslips were washed three times with PBS containing 0.05% (w/v) saponin for 5 min each time. Nuclei and actin cytoskeleton were stained with Hoechst (Sigma) and rhodamine-phalloidin (Molecular Probes), respectively. Fixed samples were viewed on a Nikon A1R confocal microscope. Images were analyzed using ImageJ (NIH).

### Mice infection

C57BL/6J wild-type from the Jackson Lab (6–8 weeks old) were bred under specific pathogen-free conditions. For oral infections, water and food were withdrawn 4 h before per os (p.o.) treatment with 20 mg/100 μL streptomycin per mouse. Afterward, animals were supplied with water and food ad libitum. At 20 h after streptomycin treatment, water and food were withdrawn again for 4 h before the mice were orally infected with 2.5 × 107 CFU/g of EIB202 or Δ*eseJ* suspension in 200 μL PBS, or treated with sterile PBS (control). Thereafter, drinking water ad libitum was offered immediately and food 2 h post-infection. At the indicated time points, mice were sacrificed and the tissue samples from the intestinal tracts, kidneys, spleens, and livers were removed for analyses.

### Zebrafish infection

Three-month old adult zebrafish (about 0.4 g) were randomly divided into groups (n=35) and infected via intramuscular injection with bacterial sample (50 cfu/fish) or PBS as a control. Fish mortality was recorded in each infection group over a period of 4 days. For immersion infection of zebrafish larvae, larvae at 5 days post-fertilization were randomly divided and immersed in PBS or PBS containing 10^5^ cfu/ml *E. piscicida* wild-type or Δ*eseJ* for 2 h. Subsequently, they were transferred to 10-cm dishes, with approximately 50 larvae in 15 ml of E3 medium per dish, and incubated at 28 °C. The bacterial colonization of every 5 fishes were then analyzed at different time points. All animal experiments were approved by the Institutional Animal Care and Use Committee of East China University of Science and Technology.

### Statistical analysis

All experiments were performed three times (as indicated in the figure legends). Statistical analyses were performed by using the student’s t-test in the SPSS software (Version 11.5, SPSS Inc.). In all cases, the significance level was defined as * *p* ≤ 0.05,** *p* ≤ 0.01 and *** *p* ≤ 0.001.

## Supporting Figures Legend

**S1 Fig.** *E. piscicida* resides and replicates in special pathogen-contained vacuoles. (A) Representative electron micrographs of *E. piscicida*-infected HeLa cells. The vocuole membranes are indicated by the white arrows. Star means *E. piscicida*. Bar = 5 μm. (B) Quantifications of galectin3 specks in HeLa cells infected with wild type EIB202 for the indicated times. Over 30 cells were analyzed for each condition. Values are means± SD (*n*= 3).

**S2 Fig.** *E. piscicida* utilizes effector EseJ to prevent lysosome maturation and fusion. (A)Representative confocal micrograph of Rab7-expressed HeLa cells after infection with *E. piscicida* EIB202 or Δ*eseJ* at an MOI of 100 for 4 h. (B) Representative confocal micrograph of ECVs colocalization with lamp-1 after infection with *E. piscicida* EIB202 or Δ*eseJ* at the indicated gentamicin incubation times. Scale bars, 20 μm. (C) Representative confocal micrograph of HeLa cells infected with GFP-labeled *E. piscicida* EIB202, ΔT3SS or Δ*eseJ* for 1 h and incubated with 100 μg/ml gentamicin for 8 h. Cells were stained with DQ Red BSA (0.25 mg/ml) for 1 h. DNA was stained using DAPI (blue).

